# Gut microbiome recovery after antibiotic usage is mediated by specific bacterial species

**DOI:** 10.1101/350470

**Authors:** Tannistha Nandi, Ivor Russel Lee, Tarini Ghosh, Amanda Hui Qi Ng, Kern Rei Chng, Chenhao Li, Yi Han Tan, David Lye, Timothy Barkham, Swaine Chen, Yunn-Hwen Gan, Barnaby Young, Niranjan Nagarajan

## Abstract

Dysbiosis in the gut microbiome due to antibiotic usage can persist for extended periods of time, impacting host health and increasing the risk for pathogen colonization. The specific factors associated with variability in gut microbiome recovery remain unknown. Using data from 4 different cohorts in 3 continents comprising >500 microbiome profiles from 117 subjects, we identified 20 bacterial species exhibiting robust association with gut microbiome recovery post antibiotic therapy. Functional and growth analysis showed that microbiome recovery is supported by enrichment in carbohydrate degradation and energy production capabilities. Association rule mining on 782 microbiome profiles from the MEDUSA database enabled reconstruction of the gut microbial ‘food-web’, identifying many recovery-associated bacteria (RABs) as primary colonizing species, with the ability to use both host and diet-derived energy sources, and to break down complex carbohydrates to support the growth of other bacteria. Experiments in a mouse model recapitulated the ability of RABs (*Bacteroides thetaiotamicron and Bifidobacterium adolescentis*) to promote microbiome recovery with synergistic effects, providing a two orders of magnitude boost to microbial abundance in early time-points and faster maturation of microbial diversity. The identification of specific microbial factors promoting microbiome recovery opens up opportunities for rationally fine-tuning pre- and probiotic formulations that prevent pathogen colonization and promote gut health.

## Introduction

The human gut microbiome harbors trillions of bacteria providing diverse metabolic capabilities and with essential roles in host health, particularly energy metabolism, immune homeostasis, and xenobiotic metabolism^1^. A stable consortium of commensal microbiota is also believed to play a key role in resisting colonization by pathogens, with dysbiosis being associated with increased risk for infections^2,3^. Several recent studies have further highlighted the importance of the gut microbiome for host health, particularly in infants and the elderly, with loss of diversity and dysbiosis being associated with various metabolic, immunological and neurological diseases^4^, and poorer response to cancer immunotherapy^5,6^.

Among the factors that can perturb the gut microbiome, antibiotic usage is known to be a major one that can cause profound and long-term alterations^7-9^. As antibiotics are widely used in healthcare, their impact on host health through microbiome dysbiosis is likely to be significant and has not been fully quantified till date^10,11^. In terms of acute response, antibiotic associated diarrhea is a common complication, while in the medium term recovery of the microbial community can be slow and variable^7-9^ and conditional on the initial state^12^. Antibiotic use can also select for drug resistance genes and organisms, thus creating a reservoir for onward transmission of resistance cassettes^13,14^. Epidemiological and model organism studies suggest that long-term consequences of antibiotic usage include immunological diseases in children^15^, metabolic diseases in adults^16^, and increased risk for infections (e.g. by *Clostridium difficile*^17^) in the elderly^18^.

Despite the mounting evidence on the importance of gut microbiome function and how antibiotic usage can severely impact it, we still do not know what enables microbiome recovery. In particular, we do not know whether specific groups of microbial taxa and the functions they perform accelerate or impede recovery and explain the variability that is seen across individuals^7-9,12^. Ecological interactions are known to play a key role in the recovery of many ecosystems^19,20^ but an analogous understanding is not currently available for the gut microbiome. A systems-level comprehension of the processes underlying gut microbiome recovery could thus aid in the design of rational interventions that reduce side-effects of antibiotic treatment and promote host health.

## RESULTS

### Identifying a microbial signature associated with gut microbiome recovery post antibiotic treatment

In order to identify microbial markers associated with gut microbiome recovery, we assembled and systematically analyzed data from 4 different cohorts (a total of 117 individuals with >500 samples). These cohorts represent individuals from 4 different countries on 3 continents (Singapore, Canada^12^, England^8^, Sweden^8^), a wide range of age groups (21-81) and using different classes of antibiotics, allowing us to infer unifying factors promoting recovery (**Table 1**). One of the cohorts is new and has not been previously analyzed (deep shotgun metagenomic sequencing of 74 samples, with >80 million reads on average), involving mostly elderly subjects from Singapore receiving inpatient antibiotic treatment (manuscript in preparation; **Suppl. Data File 1**). Overall, the diversity of the assembled cohorts enabled robustness in the analysis (via cross-validation; see **Methods**) and ensured generality of results. In addition, each cohort was analyzed independently to account for cohort-specific biases, and the results were aggregated through meta-analysis.

**Table 1.**
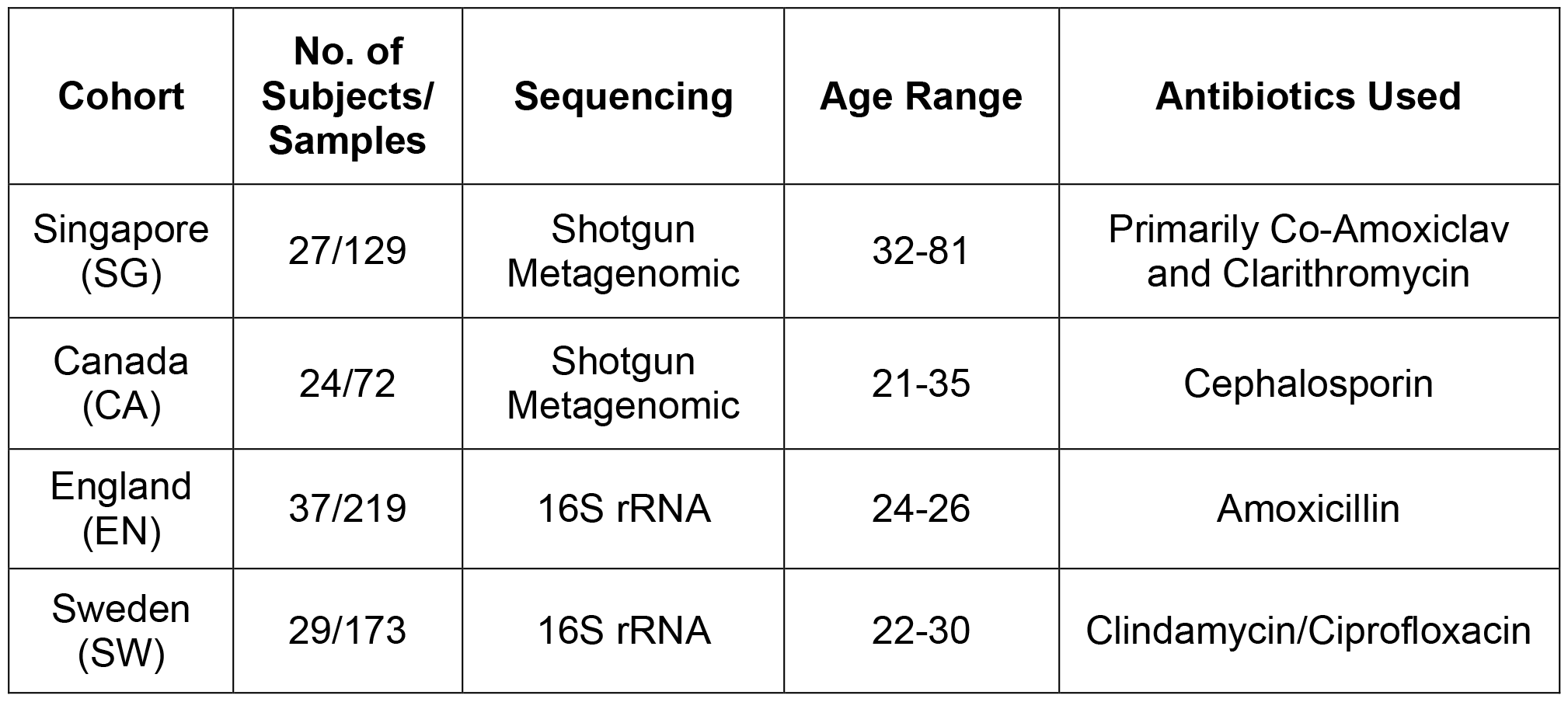
**Details of the different cohorts used in this study.**

| **Cohort** | **No. of Subjects/**<br>**Samples** | **Sequencing** | **Age Range** | **Antibiotics Used** |
| --- | --- | --- | --- | --- |
| Singapore (SG) | 27/129 | Shotgun Metagenomic | 32-81 | Primarily Co-Amoxiclav and Clarithromycin |
| Canada (CA) | 24/72 | Shotgun Metagenomic | 21-35 | Cephalosporin |
| England (EN) | 37/219 | 16S rRNA | 24-26 | Amoxicillin |
| Sweden (SW) | 29/173 | 16S rRNA | 22-30 | Clindamycin/Ciprofloxacin |

Metagenomic data from each cohort was systematically re-processed using appropriate analysis pipelines (16S rRNA or Shotgun metagenomic sequencing; **Methods**). For uniform analysis and to get balanced groups, subjects were stratified within each cohort based on median taxonomic diversity of the microbial community post antibiotics into ‘recoverers’ and ‘non-recoverers’. Recoverers exhibited a U-shaped recovery profile for gut microbial diversity, while non-recoverers start with a slightly lower median initial diversity and have even further reduced diversity up to 3 months post antibiotics (**Fig. 1A**). As expected, post-antibiotic microbiomes for recoverers were found to be more similar to pre-antibiotic microbiomes compared to non-recoverers, whose microbiomes generally appear to be diverged from unperturbed communities and dysbiotic (Mann-Whitney test *p*-value < 1.4×10^−12^; **Fig. 1B, C**). This pattern was seen to be consistent across cohorts and using different diversity metrics (**Suppl. Fig. S1**). In agreement with the notion that recovery of microbiome diversity is beneficial to the host, we also noted an enrichment of species that are known to protect against pathogen colonization in the post-antibiotic gut microbiomes of recoverers versus non-recoverers (**Suppl. Note 1**, **Suppl. Tab. S1**).

**Figure 1.**
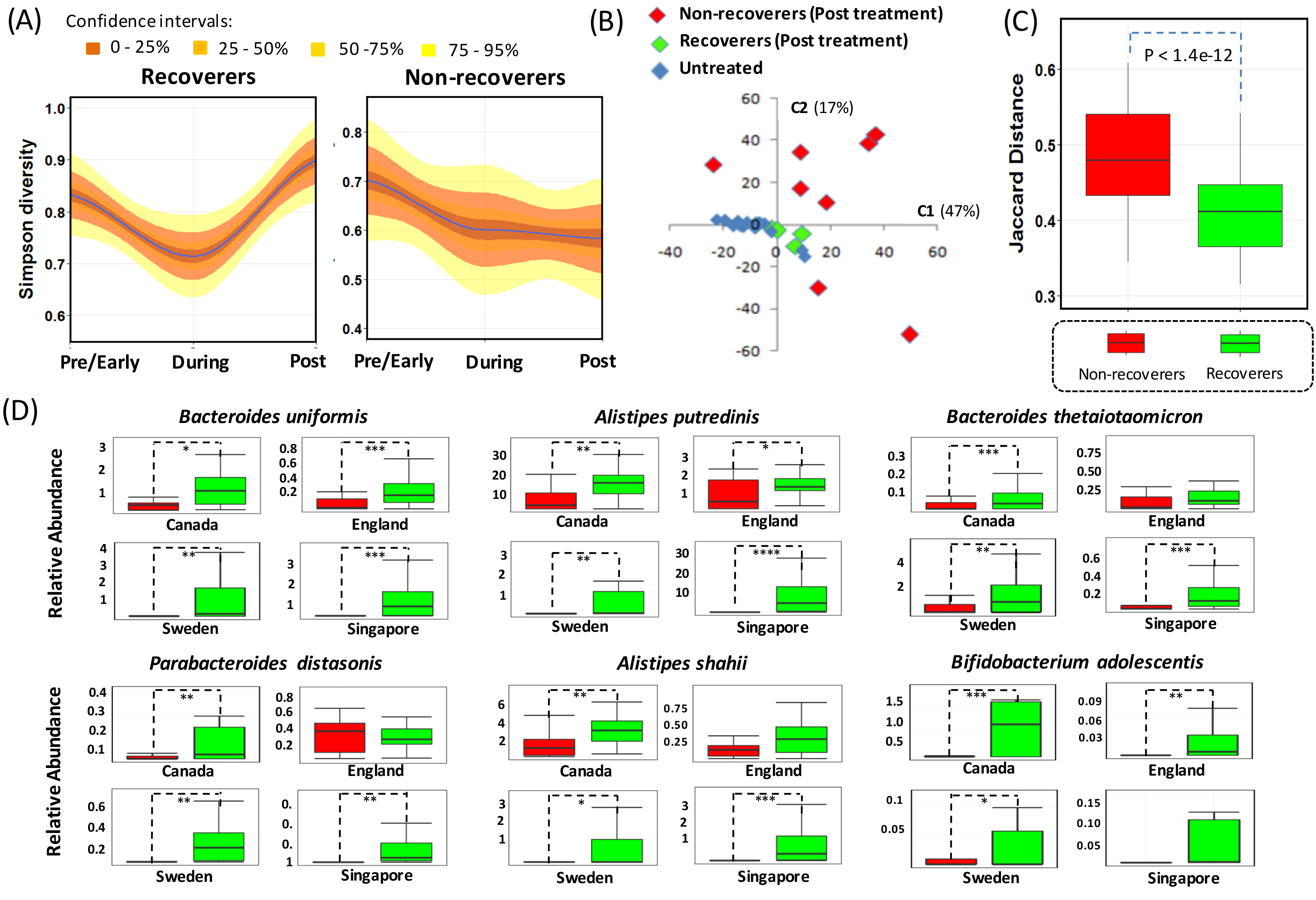
Microbial signatures associated with gut microbiome recovery. (A) Density plots showing the variation of Simpson diversities for recoverers and non-recoverers from the SG and CA cohorts across the three different stages of treatment. While the recoverers have a classic ‘U’ shaped diversity profile, non-recoverers lose diversity during antibiotic treatment and maintain that level post treatment. (B) Principal Coordinates Analysis (PCoA) plot for the CA cohort showing the dysbiosis in the post treatment gut microbiome profiles of the non-recoverers as compared to recoverers and ‘Control’ individuals (i.e. healthy volunteers who were not given any antibiotic treatment). (C) Boxplots showing the distribution of Jaccard Distances of post treatment gut microbiomes for recoverers and non-recoverers in relation to healthy controls in the ‘CA’ cohort (median value). Both (B-C) show that in addition to having higher diversity, the post treatment gut microbiome of recoverers has significantly higher similarity to microbiomes from healthy controls compared to the non-recoverers. (D) Relative abundance boxplots across cohorts for the six main species that were identified as being associated with microbiome recovery, in at least three out of four cohorts (see **Table 2** for full list). Note that ‘*’, ‘**’ and ‘***’ denote cohort-specific FDR adjusted *p*-values less than 0.1, 0.05 and 0.01 respectively.

To determine microbial taxa with a role in microbiome recovery, a two-stage approach and cross-cohort validation strategy was used to increase sensitivity and specificity of the association analysis (**Methods**; **Suppl. Data File 2**; 56 bacterial species in stage 1). Overall, 20 microbial species were identified to be significantly associated with microbiome recovery in at least two cohorts (Recovery Associated Bacteria – RAB; **Table 2**), with 6 species identified in 3 cohorts and 2 in all 4 cohorts (*Bacteroides uniformis* and *Alistipes putredinis*; **Fig. 1D**). The observed validation across diverse cohorts highlights the robustness of the associations that were observed for various taxa. In general, variability across cohorts is expected given the differences in important biological factors that could influence the gut microbiome such as diet^21^, environment^22^, genetics^23^ and the antibiotics used, though some differences could be technical as well (e.g. 16S rRNA *vs* Shotgun metagenomic sequencing). It is interesting, therefore, that despite this expected variability common associations emerge, particularly in terms of genus level homogeneity of the results (e.g. *Bacteroides* species; **Fig. 1D**; **Table 2**). We noted that many RABs have been previously shown to have beneficial impact on host health and negatively correlated with disease states, e.g. *Bacteroides uniformis* and *Parabacteroides merdae* have been observed to be negatively associated with Inflammatory bowel disease and obesity^24^, *Faecalibacterium prausnitzii* and *Roseburia inulinivorans* are known for their butyrate producing and antiinflammatory properties^24-27^, and *Bacteroides thetaiotamicron* and *Bifidobacterium* species are known for their ability to prevent pathogen colonization^27-29^ (**Table 2**). However, not all RABs are as well characterized for their contribution to gut health and importantly, their role in resilience and recovery of the gut microbiome after antibiotic treatment remains unknown.

**Table 2.**
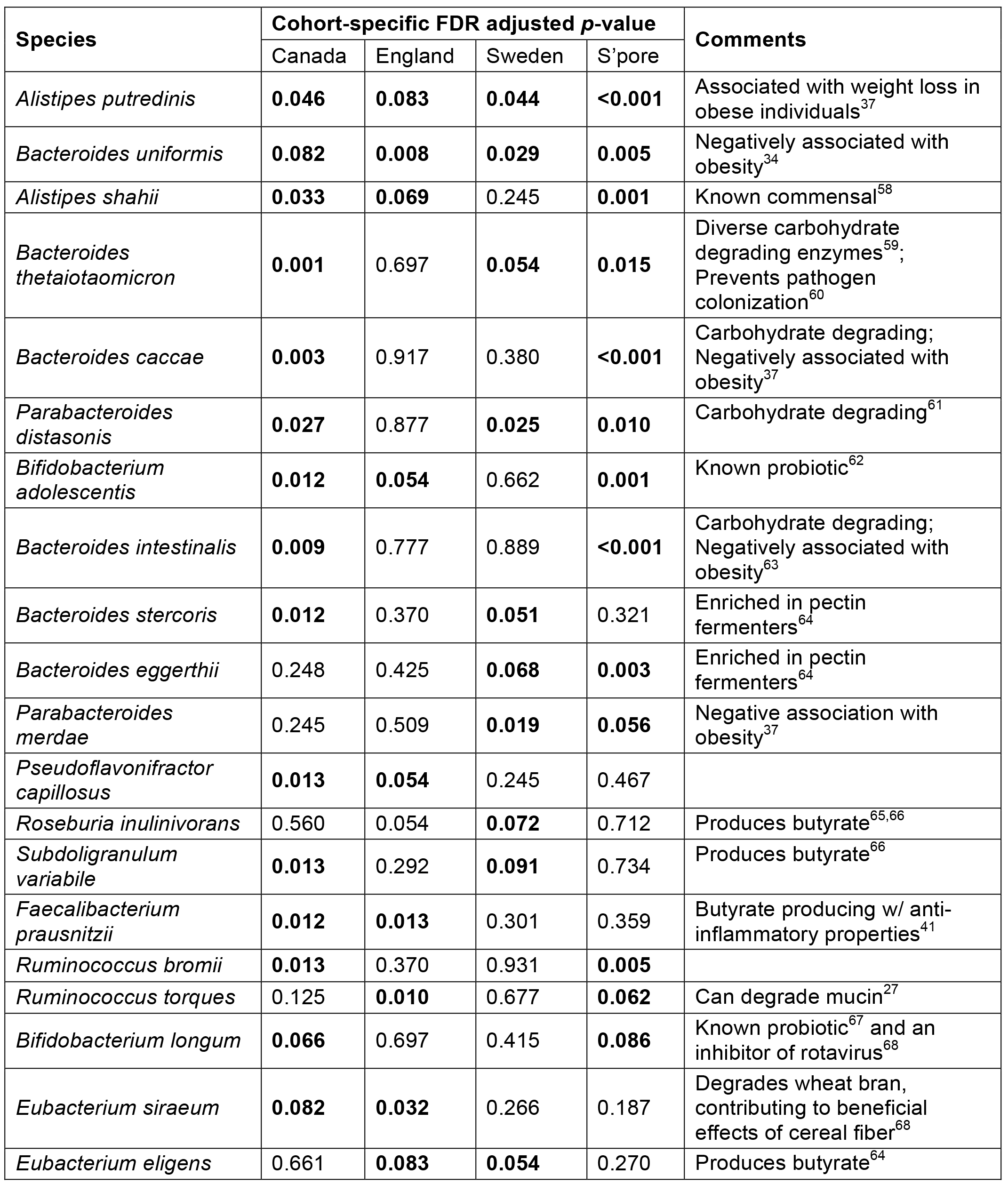
List of recovery associated bacterial taxa (RABs). RABs are ordered by the number of cohorts in which they are significantly associated (bold p-values).

We further investigated the abundance patterns of RABs across various treatment stages (pre-, during and post- antibiotics; **Methods**) and noted that while most were enriched before treatment (in recoverers *vs* non-recoverers; e.g. *Bacteroides uniformis, Bacteroides thetaiotamicron, Alistipes putredinis* and *Parabacteroides distasonis*), some were enriched in later timepoints, indicating that they may play a secondary/synergistic role in recovery as explored further later in the manuscript (**Suppl. Fig. S2**; e.g. *Bifidobacterium adolescentis* and *Faecalibacterium prausnitzii*). In addition, we explored the use of machine learning models to predict post-antibiotic recovery status from pretreatment taxonomic abundances for an individual and observed that models with moderate levels of accuracy could be obtained (70.4% from leave-one-out cross validation; **Suppl. Note 2**).

### Enrichment in carbohydrate degradation and energy metabolism capabilities is associated with bacterial growth and gut microbiome recovery

To understand microbial functions and the mechanism behind microbiome recovery, we further analyzed the metagenomic datasets that were available as part of this study (CA and SG cohorts; **Table 1**). As resistance to antibiotics could facilitate recovery, as a first hypothesis, we looked at the resistomes of recoverers and non-recoverers to see if they could explain the taxonomic differences observed (**Methods**). We noted that among the RABs, while resistance genes from *Bacteroides* and *Alistipes* species were slightly enriched in the resistomes of recoverers *vs* non-recoverers (not significant; **Suppl. Fig. 3**), genes from other RABs were not enriched in the resistomes, suggesting that other microbial functions may play a complementary role for facilitating recovery. We then compared gene families and pathways in the metagenomes of recoverers and non-recoverers to more generally identify functional capacities associated with microbiome recovery (**Methods**; FDR adjusted *p*-value < 0.1 and LDA score > 1.25; **Suppl. Data File 3**), with the analysis being restricted to pre-and during stages of antibiotic treatment to enrich for functions playing a primary role in recovery. This comparison identified a core set of growth-associated pathways pertaining to the biosynthesis of amino acids, nucleotides, co-factors and cell wall constituents (**Fig. 2**). In addition, pathways involved in carbohydrate degradation and energy production were also significantly over-represented in the gut microbiomes of recoverers. This association was also validated based on inferred pathway abundances in the English and Swedish cohorts (carbohydrate and butanaote metabolism, Mann-Whitney test *p*-value < 0.03 and 0.02 respectively; **Suppl. Fig. 4**; **Suppl. Data File 3**).

**Figure 2.**
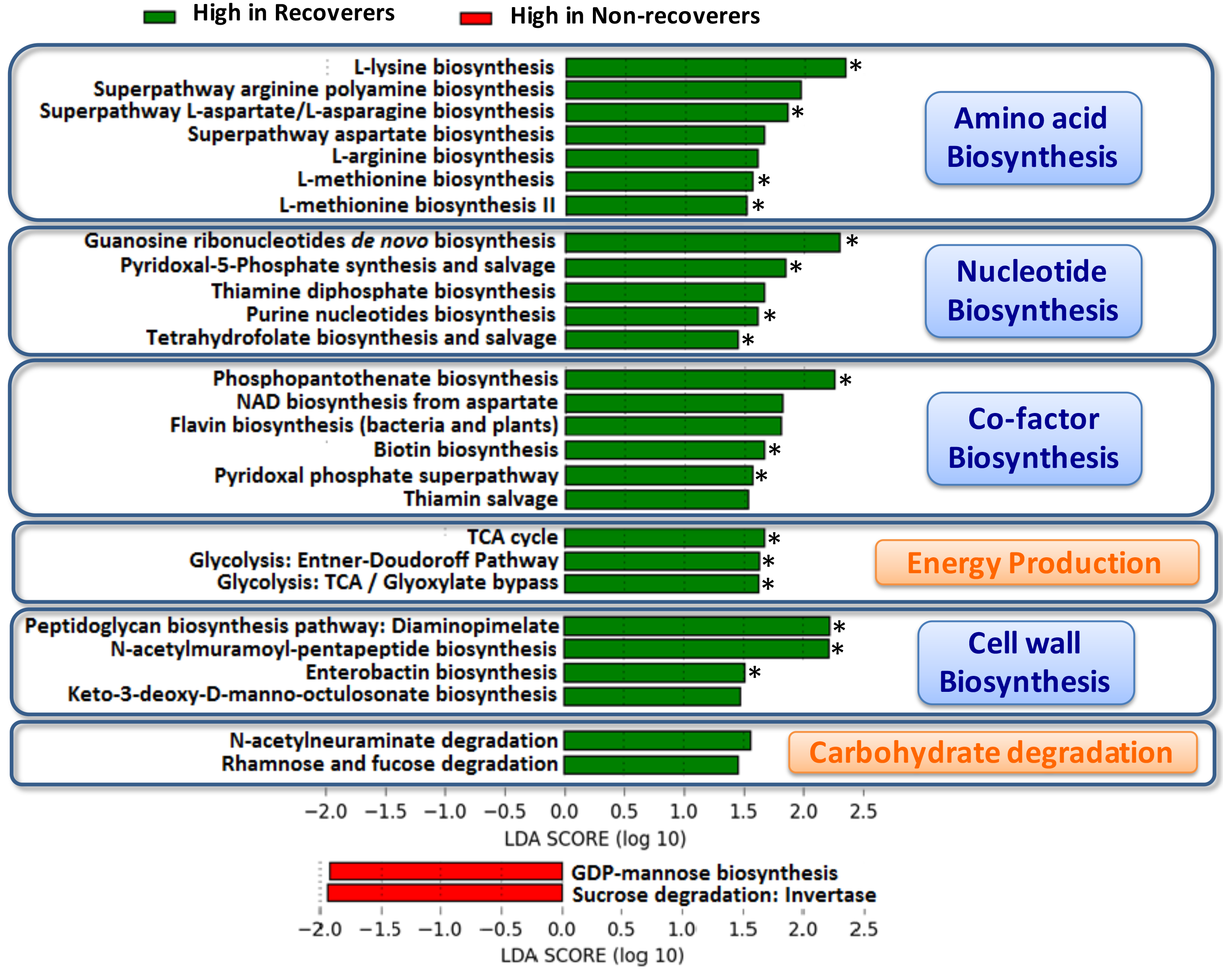
Gut microbiome functions involved in post-antibiotic recovery. Functional pathways enriched in the gut microbiomes of recoverers or non-recoverers (of the SG cohort) in the ‘Pre/Early’ and ‘During’ stages of antibiotic treatment. Note that a star (‘*’) indicates those pathways for which significant differences were also obtained in the CA cohort. Pathways were grouped into those important for energy production (in orange) and those involved in biosynthesis (in blue), highlighting the role of these two processes in microbiome recovery.

The CAZyme database of carbohydrate active enzyme families was used to further characterize the role of carbohydrate metabolism in microbiome recovery. Annotation of CAZyme families in RABs indicated that they contained a significantly higher number of CAZyme families compared to non-RABs (Mann-Whitney test *p*-value < 1e-11; **Fig. 3A**). Higher carbohydrate degrading capability thus seems to be a key shared functional property among the RABs^30^. This was also reflected at the community level where the microbiomes of recoverers was enriched in CAZyme families compared to non-recoverers (Mann-Whitney test *p*-value < 0.002 and 0.04 for CA and SG respectively; **Fig. 3B**; **Suppl. Data File 4**). The higher carbohydrate metabolism capacities of RABs could enable better nutritional harvest and thus enhance microbial growth (consistent with enriched pathways in **Fig. 2**) and subsequent recovery of gut microbiota.

**Figure 3.**
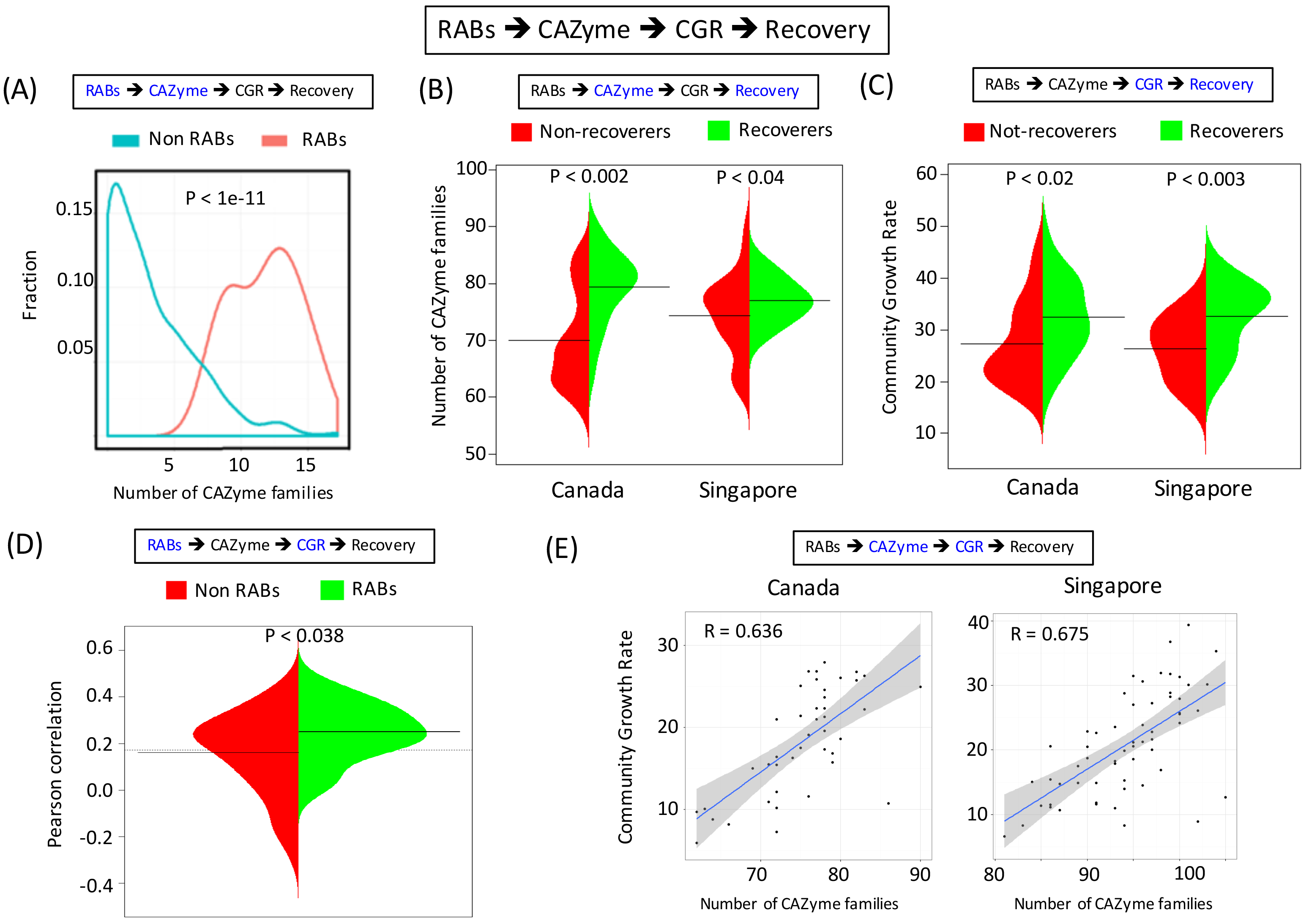
Linking microbial functions with microbiome recovery. Subfigures provide evidence for a model of microbiome recovery based on RABs being enriched for carbohydrate degradation capabilities (CAZyme), which in turn promote faster community growth (CGR), and ultimately microbiome recovery (associations shown in each subfigure are highlighted in blue). (A) Empirical distributions for the number of CAZyme families in RABs and non RABs showing that RABs are strongly enriched for CAZymes. (B) Bean plots showing the variation in the number of CAZyme families (empirical distributions) detected in the gut microbiomes of recoverers and non-recoverers in the CA and SG cohorts. In both cohorts, recoverers have more CAZyme families represented in their metagenomes (lines indicate median values; Mann-Whitney test). (C) Bean plots showing variation in the gut microbial community growth rate (empirical distributions) of recoverers and non-recoverers in the CA and SG cohorts. In both cohorts, recoverers have higher community growth rates (lines indicate median values; Mann-Whitney test). (D) Bean plot showing correlation of median abundances of RABs and non RABs in the pre-and during phase of antibiotic treatment to the post-treatment community growth rate of individuals in the SG and CA cohorts (empirical distributions). In general, the abundance of RABs is better correlated with post-treatment community growth rate (Mann-Whitney test). (E) Correlation between the number of CAZyme families detected and the overall community growth rate across all gut microbiomes constituting the CA and SG cohorts. In both cohorts, community growth rate are consistently correlated with CAZyme diversity.

To study this further from a mechanistic standpoint, we hypothesized that microbial community growth could be the intermediate phenotype that explains the association between carbohydrate active enzymes and microbiome recovery. An approach based on increased DNA around the origin of replication in replicating cells was used to infer bacterial growth rates from metagenomic data^31^ (**Suppl. Data File 5**). Using the median species-specific growth rate as a measure of community growth rate in a microbiome, we observed that recoverers exhibited higher community growth rate overall than non-recoverers across all stages of antibiotic treatment (Mann-Whitney test p-value < 0.02 and 0.003 for CA and SG cohorts, respectively; **Fig. 3C**). Additionally, we noted that the pre-and during treatment abundance of RABs had a significantly higher correlation with post-treatment community growth rate across individuals (Mann-Whitney test *p*-value < 0.038, combining CA and SG cohorts; **Fig. 3D**). Finally, consistently in both the CA and SG cohorts, community growth rate at all time points was positively correlated with the number of CAZyme families (for SG, R=0.675; for CA, R=0.636; **Fig. 3E**). Taken together, this data links the greater carbohydrate degrading potential in recovery-associated bacteria with higher microbial community growth rate and subsequent enhanced recovery of the microbiome.

### Capability to degrade host and diet-derived carbohydrates links RABs to the recovery of the microbial food web

Carbohydrate active enzymes can be varied in their function and it is possible that RABs use different combinations of functions to support microbiome recovery. To understand the role of RABs and their specific carbohydrate degradation capabilities, we took a set of 125 bacterial genomes that have been annotated for their CAZyme repertoire^30^ and clustered them based on their genome-wide profile of substrate-specific enzyme copy numbers (**Suppl. Fig. S5**). Based on this analysis, RABs were primarily observed to aggregate in 2 out of the 5 clusters obtained, with significant enrichment in cluster 1 containing genomes abundant in host (mucins) as well as diet-derived (plant and animal) carbohydrate degrading enzymes (Fisher’s exact test *p*-value < 0.038). The ability to degrade mucins is believed to play an important role in bacterial colonization of the intestine^32^. While some RABs fall in cluster 2 with genomes abundant in primarily diet-derived (plant and animal) carbohydrate degrading enzymes, only 1 RAB belongs to cluster 3 (Starch/Glycogen degradation) and no RABs were found in clusters 4 and 5 (Fungal carbohydrate and Peptidoglycan degradation), indicating that only specific carbohydrate degrading processes play a role in microbiome recovery.

Ecological interactions often play a key role in the recovery of an ecosystem^19,20^. To further study the inter-relationships between the various RABs, we sought to reconstruct a microbial ‘food-web’ that captures dependency relationships between bacteria in the gut microbiome. Association rule mining is a commonly used data mining technique to infer dependency relationships and we introduce its use here in the microbiome context. We analyzed a large database of microbiome profiles (782 samples; MEDUSA^33^; **Methods**) to identify directed binary relationships, where species A is needed for the presence of species B. The resulting network contains 192 bacterial species with 610 non-redundant edges including interactions such as the known relationships between *Bacteroides uniformis* and other *Bacteroides* and group *C. coccoides* species^34^ (**Suppl. Data File 6**).

We further investigated whether, as with most natural ecosystems, the gut microbial community could be represented by a pyramidal web, where the presence of a large number of species could be influenced by a few primary colonizers. One of the goals in this analysis was to check whether a subset of the RABs acted as primary colonizers facilitating the growth and recovery of the other species constituting the microbiome. We organized the ‘food-web’ network in terms of taxa that primarily have outgoing edges and thus support the growth of other bacteria (primary colonizers), those that have both outgoing and incoming edges (secondary colonizers), and those that primarily have incoming edges and thus depend on others for their growth (tertiary colonizers; **Fig. 4A**). Interestingly, RABs were present as primary and tertiary colonizers but not as secondary colonizers. In addition, the group of RABs that were primary colonizers were all from cluster 1 with mucin degrading capabilities, while cluster 2 RABs were restricted to being tertiary colonizers (plant and animal carbohydrate degrading). These observations are in agreement with our expectation that while some of the RABs may be essential to microbiome recovery and act as keystone species (primary colonizers), others may only play a supportive role or serve as indicator species for microbiome recovery (tertiary colonizers).

**Figure 4.**
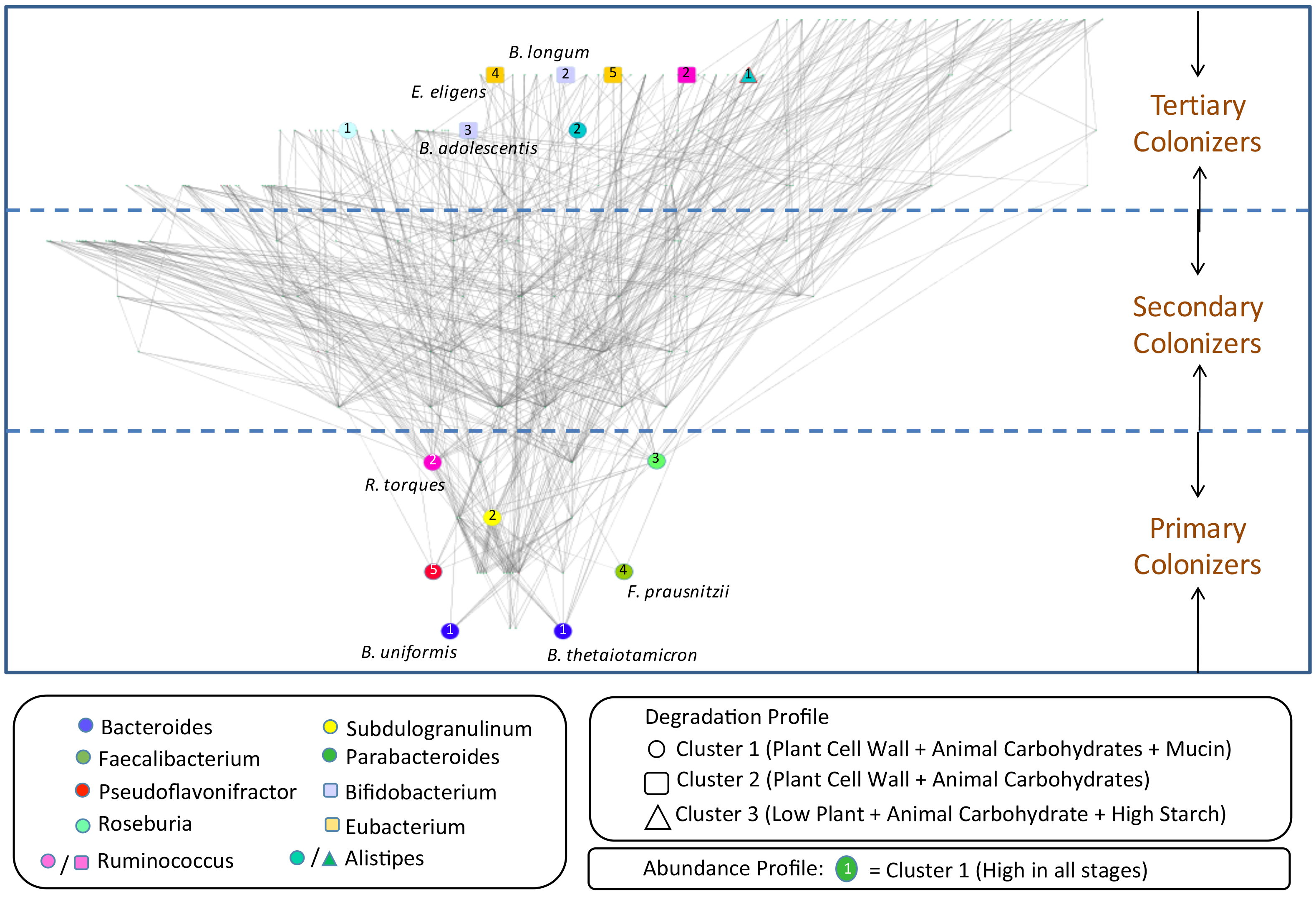
Role of RABs in recovery of the microbial food web. (A) Graph showing network structure of microbial dependencies inferred using an association rule mining approach, where an edge from species A to species B indicates that A’s presence is required to have B in the community. Nodes are ordered from the bottom to the top such that species at the bottom have more outgoing edges than incoming edges (‘Primary colonizers’), while species at the top have more incoming edges than outgoing edges (‘Tertiary colonizers’). RABs (highlighted in different colors based on the genus they belong to) were observed either at the bottom or top of the graph. RABs at the bottom of the graph were exclusively from cluster 1 (degradation profile; **Suppl. Fig. S5**), defined by mucin degrading CAZymes. Clusters based on abundance profile over time (**Suppl. Fig. S2**) are indicated using numbers and do not seem to be biased in different regions of the graph. (B) Schematic representation of the gut showing a model for microbiome recovery based on these observations. RABs from cluster 1 (**Suppl. Fig. S5**) colonize the epithelial mucosa better because of their mucin degrading capabilities (step 1), and since they can also break down dietary plant and animal derived carbohydrates (step 2), they act as primary colonizers that facilitate the growth of nonprimary colonizers (step 3). Some of these secondary and tertiary colonizers may be better adapted to degrading plant and animal carbohydrates. The overall activity of primary and non-primary colonizers results in producing simpler sugars (promoting the growth of more bacteria (step 4) and short chain fatty acids (SCFAs), which are then utilized by colonocytes for their growth leading to increased mucin production (step 5). This positive feedback loop promotes faster recovery of microbial biomass and community diversity to re-establish homeostasis in the gut.

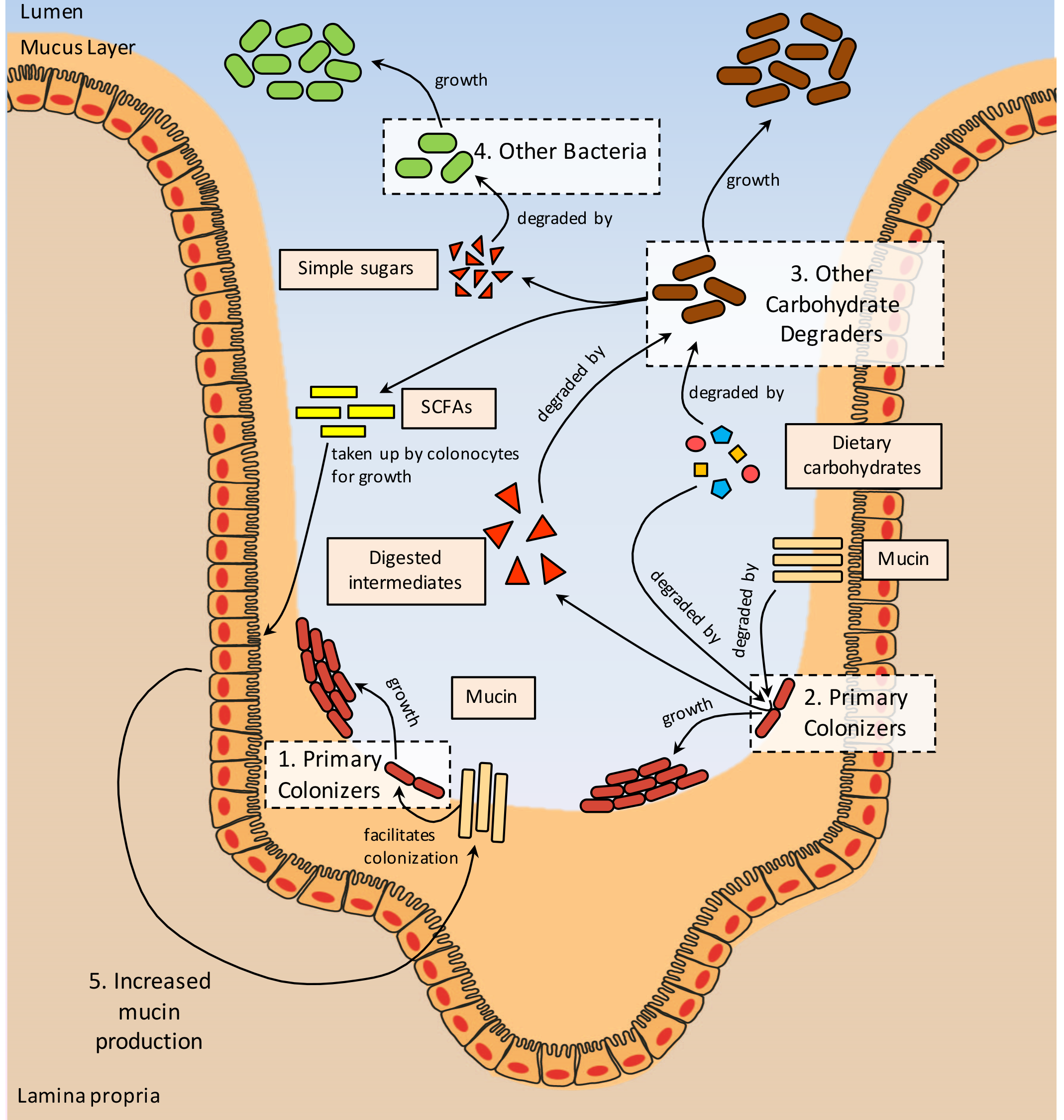

Overall, the carbohydrate degradation profiles of RABs and their inter-relationships in the food-web suggest a model for how they interact in the context of microbiome recovery (**Fig. 4B**). RABs that belong to cluster 1 can degrade mucin in addition to diet-derived carbohydrates, making them adept as primary host colonizers that can also break down complex carbohydrates for use as energy sources by other bacteria. This can facilitate the growth of non-host-colonizing, complex carbohydrate degraders in the gut, as well as other bacteria that rely on the simple sugars produced for their growth. Furthermore, production of SCFAs (particularly butyrate) by RABs and the recovering bacterial community can stimulate mucin production by colonocytes providing a positive feedback loop that can contribute to accelerated microbiome recovery^35,36^.

### Primary and tertiary colonizing RABs synergistically enhance microbiome recovery *in vivo*

Microbiome recovery is likely to be a multi-stage process involving several bacteria with different roles in different individuals. To begin to understand these interactions based on the RABs identified in this study, we conducted a proof-of-concept experiment in a mouse model to see if some of our human observations could also be qualitatively recapitulated in mice. Specifically, we gave mice antibiotics for 5 days, followed by oral gavage of different RABs (*B. thetaiotamicron* - Bt, *B. adolescentis* - Ba), negative controls *(Bacillus spp.* - Bsp, PBS) and combinations *(B. thetaiotamicron* and *B. adolescentis* - Bt+Ba, *Bacillus spp.* and *B. adolescentis* - Bsp+Ba; **Methods**). Stool samples were then collected every three days for up to 22 days and analyzed using shotgun metagenomic sequencing (**Methods**; **Fig. 5A**). In total, the study involved 6 groups, with 2-6 cages per group (each cage with 2 mice) and 9 timepoints (243 metagenomic libraries).

**Figure 5.**
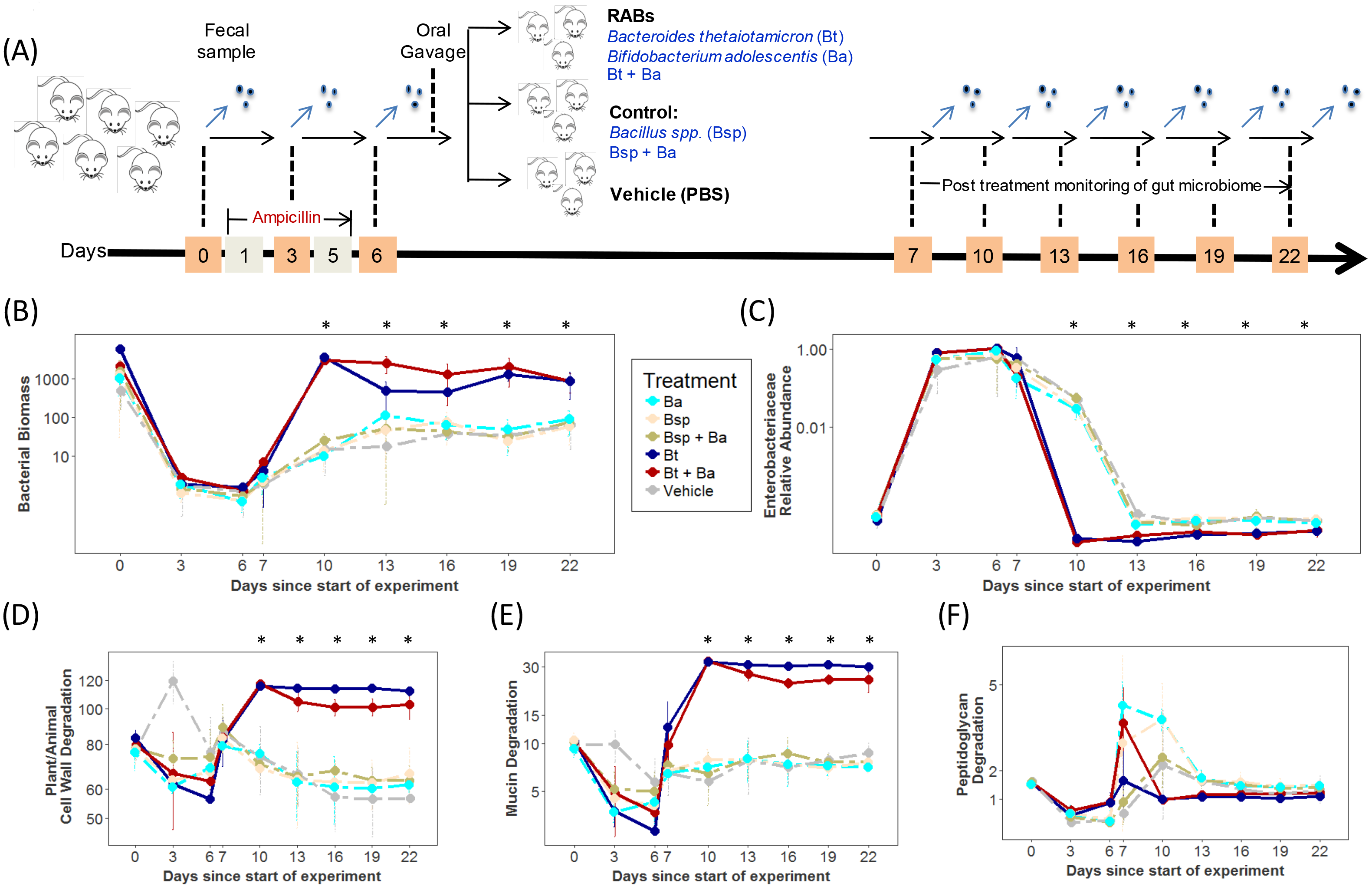
Promoting microbiome recovery in a mouse model using RABs. (A) Schematic depicting the design of a mouse model experiment to study the impact of RABs in promoting microbiome recovery. Mice were given antibiotics for 5 days, followed by a rest day and gavage of different RABs and controls. Shotgun metagenomics was then used to monitor microbiome changes every 3 days. (B) Microbial biomass (median ± 1 s.d.) in different groups of mice across time (excluding gavaged species). Stars in all subfigures (‘*’) indicate timepoints where the Bt and Bt+Ba groups were significantly different from other groups (Mann-Whitney test *p*-value < 0.05). (C) Relative abundance of Enterobacteriaceae (median ± 1 s.d.) in different groups of mice across time. (D, E, F) Reads per million (RPM) mapping to CAZymes associated with plant/animal cell wall, mucin and peptidoglycan degradation, respectively, across different experimental groups and timepoints.

Overall, all treatment groups exhibited a >3-log reduction in microbial biomass after antibiotic treatment as expected (**Methods**; **Fig. 5B**). However, starting from 1 day after gavage (day 7), and more noticeably 4 days after gavage (day 10), the Bt and Bt+Ba groups exhibited similarly enhanced recovery (>100×) of microbial biomass compared to other groups (**Fig. 5B**; **Suppl. Fig. S6A**). This is not explained by colonization of the gavaged species alone, as reads belonging to them were removed before doing this analysis. Interestingly, despite being a known probiotic, gavage with Ba alone did not promote enhanced recovery of biomass, while Bt+Ba supported more stable recovery compared to Bt alone (**Fig. 5B**; **Suppl. Fig. S6B**). While the Bt and Bt+Ba groups converge to their microbial biomass at pre-antibiotic levels by day 10, all other groups continue to have lower biomass at day 22. Similar trends were also seen in terms of microbial community profiles, with the Bt and Bt+Ba groups being more similar to the diversity of the pre-antibiotic microbiome at day 10 than in other groups (**Suppl. Fig. S6B**). However, the Bt+Ba group appears to be better in recovering a pre-antibiotic microbiome at day 22 compared to Bt alone, indicating that synergistic interactions between Bt and Ba may play a role here.

The Bt and Bt+Ba groups also exhibited lower levels of Enterobacteriaceae compared to other groups from day 10 onwards (**Fig. 5C**), and similar patterns were seen for other potential pathogens as well (e.g. *Clostridium difficile, Chlamydia trachomatis* and *Staphylococcus aureus*). These observations are suggestive of a form of colonization resistance that may be provided by the enhanced microbiome recovery in these two groups. Recapitulating the observations in human cohorts, we also observed enhanced relative abundance of enzymes for plant and animal cell wall degradation as well as mucin degradation in the groups exhibiting faster recovery (**Fig. 5D, E**). As a control comparison, peptidoglycan degrading enzymes were not specifically enriched in the Bt and Bt+Ba groups, in agreement with our earlier observation that these functions are not defining characteristics for RABs (**Fig. 5F**, **Suppl. Fig. S5**).

## Discussion

The bacterial species and functions identified in this study provide a first, data-driven view of how shared microbial factors contribute to gut microbiome recovery in diverse human cohorts around the world. Our findings emphasize the central role of enabling energy harvest from diet and the ability to colonize the host in the keystone species that underpin ecological recovery, while antibiotic resistance in general plays a less important role. As environmental factors strongly influence the gut microbiome^22^, the specific keystone species that are important for an individual could additionally vary with host and dietary factors. Uncovering these in larger cohorts should be feasible using similar analytical approaches as used here, and could help train antibiotic and environment-specific machine learning models to predict microbiome recovery. Such models would have clinical utility, especially for at-risk elderly or cancer patients, to guide targeted intervention and prevention strategies.

Consistent with our emerging understanding of how diet modulates the gut microbiome^21,22^, another perspective from which to see our results is the importance of feeding gut bacteria correctly (in addition to having the right species) to promote recovery. Many of the identified RABs are specialist carbohydrate fermenters (e.g. pectin) and a high fiber/low fat diet could aid in selecting and expanding them. For example, in a study on how gut microbiota differ in twins discordant for obesity control metabolism, Ridaura *et al* identified 4 RABs (*B. uniformis, B. thetaiotaomicron, Alistipes putredinis* and *Parabacteriodes merdae*) as being transplantable features of a “lean microbiome”, but transplantation was dependent on a high fiber diet^37^. Similarly, pectin supplementation can promote species from the Bacteroidetes phylum with associated improvement in gut barrier function^38^, as well as more stable fecal microbiota transplantation^39^. Finally, different oligosaccharides can promote the growth of several butyrate producing RABs^40,41^ (**Table 1**), contributing to microbiome recovery by reducing host inflammation and increasing mucin production^36^.

In general, ecological theory has suggested that ecosystem recovery is a complex, multi-step process that is determined by interactions between many species^19,20^. Similar properties are likely to hold for the human gut microbiome with multiple RABs and the synergistic interactions between them playing a role to promote microbial cross-feeding, enable biomass recovery, modulate host inflammation, prevent pathogen colonization and eventually regain taxonomic and functional diversity. While results from our mouse model have provided initial hints, further exploring combinatorial interactions between RABs is likely more feasible using *in vitro* co-culture^42^ or *in silico* metabolic models^43,44^. Metabolic modeling could, in particular, help explore the contributions of different carbohydrate degradation genes and processes to microbiome recovery^44^, especially for many anaerobic bacteria that are hard to culture or genetically modify^45^. Such investigations could also be informative in understanding the contributions of core and accessory genomes within a species and whether strain-level differences could cause variability in microbiome recovery across individuals.

Conceptually, the microbial ‘food-web’ as data-mined in this study is a powerful resource for organizing our understanding of how microbes interact and assemble in the human gut. By using a large database of human gut microbiome profiles, we can determine microbial assemblages that are feasible and the dependency relationships that they suggest. These can then help interpret longitudinal studies of recovery and infer the succession of species that play a role. While our current work suggests that introduction of primary colonizers such as *B. thetaiotamicron* may be a necessary and sufficient way to reduce dysbiosis in comparison to existing probiotics such *B. adolescentis*, synergistic combinations could provide other benefits such as colonization resistance against opportunistic pathogens. Further studies of cross-feeding interactions in the ‘food-web’ may also help identify prebiotics that could serve as supplements to accelerate the process of gut microbiome recovery. In general, understanding microbiome recovery post antibiotic treatment sets the stage for a more general understanding of how microbiome dysbiosis in other diseases could be reverted back to a healthy state using individual-specific pre-and probiotic formulations.

## METHODS

### Study Populations

*(a) Singapore*: The Singaporean cohort (‘SG’; manuscript in preparation) is a natural history cohort consisting of individuals admitted to Tan Tock Seng Hospital (TTSH) in Singapore and prescribed antibiotics for 1-2 weeks (**Table 1**). Stool samples were collected as soon as possible after admission (pre-/early: <3 days into treatment), during and up to 3 months after antibiotic usage. The study was approved by the Institutional Review Board at TTSH (DSRB 2013/00769).

*(b) Canada*: Shotgun metagenomic datasets for a Canadian cohort^12^ (‘CA’) were obtained from the European Nucleotide Archive database (Study Accession Number: PRJEB8094; **Table 1**). The study analyzed fecal samples from healthy individuals who were administered antibiotics (three timepoints: pre-antibiotic day 0, during treatment day 7 and post treatment day 90).

*(c) England and Sweden*: 16S rRNA sequencing datasets for an English and a Swedish cohort^8^ (‘EN’, ‘SW’) were obtained from the NCBI short read archive (Project ID: SRP057504; **Table 1**). In both cohorts, healthy volunteers were given antibiotics and fecal samples analyzed for day 0 (pre-antibiotic), day 7 (during treatment) and for one and two month follow-ups (post treatment).

For the CA, EN and SW cohorts, all antibiotic treated subjects with data from the 3 treatment stages were further analyzed to identify recovery associated bacterial taxa and functions.

### DNA extraction and sequencing for SG cohort

Extraction of DNA from stool samples was carried out using PowerSoil DNA Isolation Kit (MoBio Laboratories, California, USA) with minor modifications to the manufacturer’s protocol (volume of solutions C2, C3 and C4 were doubled and centrifugation time was extended to twice the original duration). Purified DNA was eluted in 80μl of Solution C6. DNA libraries were prepared by using 20ng of extracted DNA re-suspended in a volume of 50μl and subjected to shearing using Adaptive Focused Acoustics™ (Covaris, Massachusetts, USA) with the following parameters; Duty Factor: 30%, Peak Incident Power (PIP): 450, 200 cycles per burst, Treatment Time: 240s. Sheared DNA was cleaned up with 1.5× Agencourt AMPure XP beads (A63882, Beckman Coulter, California, USA). End-repair, A-addition and adapter ligation was carried out using the Gene Read DNA Library I Core Kit (Qiagen, Hilden, Germany) according to the manufacturer’s protocol. Custom barcode adapters (**Suppl. Table 1**) were used in place of GeneRead Adapter I Set for adapter ligation. DNA libraries were cleaned up twice using 1.5× Agencourt AMPure XP beads (A63882, Beckman Coulter, California, USA) before enrichment of libraries using the protocol adapted from Multiplexing Sample Preparation Oligonucleotide kit (Illumina, California, USA). Enrichment PCR was carried out with PE 1.0 and custom index-primers (**Suppl. Table 1**) for 14 cycles. Libraries were quantified using Agilent Bioanalyzer and prepared with Agilent DNA1000 Kit (Agilent Technologies, California, USA), pooled in equimolar concentrations. Sequencing of the samples was performed using the Illumina HiSeq 2500 (Illumina, California, USA) sequencing instrument to generate >80 million 2×101 bp reads on average.

### Taxonomic and functional profiling for all cohorts

For metagenomic sequencing datasets (CA and SG cohorts) raw reads were quality filtered and trimmed using default options in famas (https://github.com/andreas-wilm/famas). Reads that are potentially from human DNA were removed by mapping to the hg19 reference using BWA-MEM^46^ (default parameters; coverage >80% of read). The remaining reads were used for taxonomic profiling using MetaPhlAn with default parameters^25,47^ (**Suppl. Data File 1**). Functional profiles for the metagenomes were obtained using the HUMAnN2 program [14] (**Suppl. Data File 3**).

For the 16S rRNA sequencing datasets (EN and SW cohorts) taxonomic classification was done by mapping reads to the SILVA database^48^ (v123), using BLASTn [7]. For each read, the species corresponding to the best hit (with identity > 97% and query coverage > 95%) was obtained and was taken as the source species of the read. In the case of multiple hits, the source taxon was computed as the Lowest Common Ancestor of the hit species. Reads assigned to each taxon were aggregated to obtain a relative abundance profile for each sample (**Suppl. Data File 1**). PICRUSt^49^ was used to infer KEGG pathway abundances from the corresponding taxonomic profiles (**Suppl. Data File 3**).

### Identification of recovery associated bacterial taxa and functions

Individuals were classified as ‘recoverers’ and ‘non-recoverers’ in each cohort to enable cohort-specific association analysis and identification of recovery associated bacterial taxa and functions. The median post-treatment Simpson diversity of the microbiome (species level) was used as the threshold for this classification in each cohort to provide balanced groups. Samples within a 10% window of the interquartile range from the median were marked as having indeterminate status and excluded from further analysis. A classification approach rather than regression analysis was used as the observed diversity values were not well distributed across the range of values. A two-stage approach was used to combine results from all cohorts to sensitively identify recovery associated taxa and an across-cohort validation strategy was used to identify taxa that are significant in at least 2 out of 4 cohorts. In stage 1, a non-parametric test was used within each cohort (Mann-Whitney test) to filter candidate taxa (*p*-value > 0.05). The remaining candidates were merged across cohorts to compute a combined *p*-value using Fisher’s method and filtered with a FDR adjusted *p*-value threshold of 0.1 (Bonferroni-Hochberg method). Next, in stage 2, cohort-specific FDR adjusted p-values were re-computed for this subset of taxa and only taxa with consistent (in terms of direction of change) significant associations (FDR < 0.1) in at least 2 cohorts were retained. This analysis was done within each treatment stage (pre-, during and post-antibiotics) as well as jointly to increase sensitivity in identifying recovery associated taxa regardless of treatment stage.

Functional profiles computed with HUMAnN2 were compared between recoverers and non-recoverers in the SG and CA cohorts using the linear discriminant analysis approach in LEfSe^50^ (version 1.1.0) to identify differentially abundant pathways.

### Microbial community growth rate analysis

An *in silico* approach, originally proposed by Korem et al^31^, was used to compute the skew of DNA copy number starting from around the origin of replication to the termination region (peak-to-trough ration or PTR), as an estimate of growth rates for individual species in the microbiome from shotgun metagenomic data (PTRC1.1: https://genie.weizmann.ac.il/software/bacgrowth.html, default parameters). The median PTR value for species in a community was then used to represent community growth rate (CGR) for each sample (**Suppl. Data File 5**).

### Profiling of carbohydrate active enzymes (CAZymes)

An in-house nucleotide gene database for CAZymes was created by downloading sequences from NCBI corresponding to Accession IDs for different CAZyme families annotated in dbCAN^51^ (http://csbl.bmb.uga.edu/dbCAN/). Metagenomic reads were mapped to this database for each sample with BWA-MEM^46^ (default parameters) to compute the fraction of reads mapping to the CAZyme gene per kbp per million reads in the metagenome (RPKM). Results were aggregated for each CAZyme family based on values for individual CAZyme genes belonging to a family.

### Analysis of antibiotic resistance genes within gut microbiomes

Resistome profiling within a microbiome was performed similarly by mapping metagenomic reads using BWA-MEM (default parameters) to the ARG-ANNOT database^52^, and calculating the fraction of reads mapping to a resistance gene per kbp per million reads of the metagenome (RPKM). Kraken^53^ was used with default parameters to obtain the taxonomic classification of reads and thus obtain the relative representation of different taxonomic groups within the resistome.

### Clustering of species based on their carbohydrate degradation profiles

The substrate-specificities of different Glycoside hydrolase (GH) and Polysaccharide lyase (PL) families was obtained from previous studies^27,28^. These included substrates such as plant cell wall carbohydrates, animal carbohydrates, peptidoglycans, fungal carbohydrates, sucrose/fructose, dextran, starch/glycogen and mucins. Copy number annotations for each GH and PL family in 125 bacterial species were obtained from a previous genome scale analysis of CAZymes in species belonging to the human gut microbiome^30^. Copy numbers of GH/PL genes within each of the 8 substrate specificities were aggregated and normalized to obtain an overall carbohydrate degradation profile for each bacterial species. Degradation profiles were then clustered using hierarchical clustering (‘hclust’ function in R with Euclidean distance and complete linkage clustering) to group species based on their enzyme repertoire for different categories of carbohydrates. Association of the identified recovery associated bacteria to one or more of these clusters was then evaluated using Fisher’s exact test.

### Construction of microbial ‘food web’ using association rule mining

To identify directed associations between bacterial species where the presence of one is important for the presence of another (but not *vice versa*), a data-mining technique called ‘association rule mining’^54^ was applied to a large public collection of gut microbiome profiles in the MEDUSA database^33^ (782 gut microbiome profiles from USA, China and Europe). To convert relative abundance profiles from MEDUSA into presence-absence profiles (1 if a species is present and 0 otherwise), relative abundances 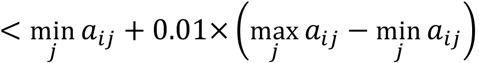, i.e. within 1% of the minimum relative abundance values *a*_*ij*_ for species *i* across subjects *j*, were assumed to be due to technical noise. Binary association rules between species were then inferred using the apriori algorithm implemented in the R package ‘arules’ (using Confidence threshold of 0.95 and Support threshold of 0.05). After removal of transitive edges and symmetric relationships, a total of 610 directed association edges remained across 192 species (**Suppl. Data File 6**). Association edges and corresponding nodes for species were plotted using the hierarchical layout of Cytoscape, where the hierarchical level of a species was based on the difference between the number of outgoing and incoming edges.

### Promoting microbiome recovery in a mouse model

#### Ethics statement

Mouse experimental protocols were reviewed, approved and carried out in strict accordance to the recommendations by the Institutional Animal Care and Use Committee from the National University of Singapore.

#### Bacterial strains and culture conditions

Lyophilized probiotic strains (ATCC 29148 *Bacteroides thetaiotaomicron*, DSM 20083 *Bifidobacterium adolescentis*) were revived in TSB media supplemented with 5% defibrinated sheep blood under anaerobic conditions at 37°C. Upon revival, *B. thetaiotaomicron* was subcultured and maintained in TYG media, whereas *B. adolescentis* and an environmental *Bacillus* isolate were subcultured and maintained in BHI media.

#### Antibiotic administration and inoculation with test strains

Eight-week-old C57BL/6J male mice from a single breeding colony were gavaged individually with 2.5 mg ampicillin per day for 5 days under specific pathogen-free conditions. Upon cessation of antibiotic treatment, mice were allowed to recover for 24 hours, before 2-6 cages of mice (two mice per cage) were each orally inoculated with: A) 5 × 10^7^ CFUs *B. thetaiotaomicron*, B) 5 × 10^7^ CFUs *Bacillus spp*., C) 5 × 10^7^ CFUs *B. adolescentis*, D) 5 × 10^7^ CFUs *B. thetaiotaomicron* + 5 × 10^7^ CFUs *B. adolescentis*, E) 5 × 10^7^ CFUs *Bacillus spp*. + 5 × 10^7^ CFUs *B. adolescentis*, or F) phosphate-buffered saline (PBS). Strains were transported from anaerobic chamber to animal facility via anaerobic “balch-type” culture tubes with aluminum seals (Chemglass Life Sciences, New Jersey, USA).

#### Fecal sample collection and DNA extraction

Fecal pellets were freshly collected as a cage unit (two mice per cage) over multiple times points: before antibiotic treatment (Day 0), mid-point of antibiotic treatment (Day 3), end-point of antibiotic treatment (Day 6), 1-day post-gavage (Day 7), 4-days post-gavage (Day 10), 7-days post-gavage (Day 13), 10-days post-gavage (Day 16), 13-days post-gavage (Day 19) and 16-days post-gavage (Day 22). Total bacterial DNA was extracted from fecal samples using the PowerSoil DNA isolation kit (MoBio Laboratories) according to the manufacturer’s instructions.

#### Library preparation and deep sequencing

Community DNA extraction was carried out using PowerSoil DNA Isolation Kit (MoBio Laboratories, California, USA) according to the manufacturer’s protocol, without modifications. Library preparation and deep sequencing was performed as described for the human fecal samples obtained from the SG cohort (described earlier under the section ‘DNA extraction and sequencing for SG cohort’) with the following modification: 50 ng of DNA was used as input for preparation of libraries.

#### Taxonomic profiling

For obtaining the taxonomic profiles of the mouse gut metagenomes, reads were mapped to the NR database using DIAMOND^55^. The taxonomic classification of each sequence was then obtained by using the LCA-based approach in MEGAN^56^ (default parameters, minimum score of 50).

#### Calculation of microbial biomass

Bacterial biomass (up to a constant factor) was estimated by taking all reads classified to bacterial taxa and normalizing by non-microbial reads. Specifically, plant or host-derived reads were used, respectively, based on the assumption that the absolute amounts of their DNA would remain roughly constant in the analyzed mouse fecal samples. Similar trends were observed for both forms of normalization (default=plant normalized), normalization based abundances were found to correlate with qPCR estimates (plant normalized, r=0.73, *p*-value=10^−4^; host normalized, r=0.82, *p*-value=3.5×10^−6^), and the observed differences between Bt and Bt+Ba groups versus other groups were also validated using qPCR (day 10, fold-change=94-170×). Note that sequencing based biomass estimates have the advantage that they allow us to subtract reads belonging to the gavaged species and are also not affected due to variations in 16S rRNA copy number across taxa.

#### qPCR Analysis

Absolute quantification of the 16S rRNA gene was done by quantitative PCR (qPCR). A pair of universal 16S bacterial primers^57^ were used to amplify DNA extracted from the six different treatment groups on days 0, 3, 10 and 13 (**Suppl. Table 1**). Reactions were prepared on a 384-well plate, in triplicates, using 5 μL of PowerUp SYBR Green Master Mix (Thermo Fisher Scientific, Massachusetts, USA), 0.5 μL of 5μM primers and 1 μL of 10× diluted DNA, in a total volume of 10 μL for each reaction. The ViiA 7 Real-Time PCR System (Thermo Fisher Scientific, Massachusetts, USA) was used for qPCR with the following amplification parameters: 1 cycle of 95°C for 2 min, 40 cycles of 95°C for 15 s, 60°C for 15 s, and 72°C for 1 min. A standard curve was created using serial dilution of synthesized double-stranded DNA oligomers (gBLOCK, Integrated DNA Technologies, Inc., Iowa, USA; **Suppl. Table 1**) to convert CT values to copy numbers. Copy numbers from day 0 were used to scale bacterial abundances to the same starting baseline.

## Data Access

Illumina sequencing data for this study (mouse models) has been deposited to the Sequence Read Archive under project ID SRP142225 (reviewer metadata link: ftp://ftp-trace.ncbi.nlm.nih.gov/sra/review/SRP142225_20180423_152835_2726e05d1c01c63b0742fdbb3d89c0bc).

## Acknowledgements

This work was supported by funding from the National Healthcare Group (NHG-CSCS/12008) and A*STAR, Singapore.

